# Photografted zwitterionic hydrogel coating durability for reduced foreign body response to cochlear implants

**DOI:** 10.1101/2022.06.24.497518

**Authors:** Adreann Peel, Douglas M. Bennion, Ryan Horne, Marlan R. Hansen, C. Allan Guymon

## Abstract

**Objective:** Durability of photografted zwitterionic hydrogel coatings on cochlear implant biomaterials was examined to determine viability of these antifouling surfaces during insertion and long-term implant usage. Approach: Tribometry was used to determine the effect of zwitterionic coatings on lubricity of surfaces with varying hydration level, applied normal force, and timeframe. Additionally, flexural resistance was investigated using mandrel bending. *Ex vivo* durability was assessed by determining coefficient of friction between tissues and treated surfaces. Furthermore, cochlear implantation force was measured using cadaveric human cochleae. Main results: Hydrated zwitterionic hydrogel coatings reduced frictional resistance approximately 20-fold compared to uncoated PDMS, which importantly led to significantly lower mean force experienced by coated cochlear implants during insertion compared to uncoated systems. Under flexural force, zwitterionic films resisted failure for up to 60 minutes of desiccation. The large increase in lubricity was maintained for 20 hours under continual force while hydrated. For loosely crosslinked systems, films remained stable and lubricious even after rehydration following complete drying. All films remained hydrated and functional under frictional force for at least 30 minutes in ambient conditions while drying, with lower crosslink densities showing the greatest longevity. Moreover, photografted zwitterionic hydrogel samples showed no evidence of degradation and nearly identical lubricity before and after implantation. Significance: This work demonstrates that photografted zwitterionic hydrogel coatings are sufficiently durable to maintain viability before, during, and after implantation. Mechanical properties, including greatly increased lubricity, are preserved after complete drying and rehydration for various applied forces. Additionally, this significantly enhanced lubricity translates to significantly decreased force during insertion of implants which should result in less trauma and scarring.

## 1. Introduction

Medical implants have advanced significantly in recent years, expanding capability and function.^1^ In particular, cochlear implants (CIs) have increasingly been used to rehabilitate hearing loss for those who suffer from severe to profound sensorineural hearing loss. Recent advancements in “hybrid” CIs enable restoration of high frequency hearing via electrical stimulation while preserving residual acoustic hearing in the lower frequencies in patients with only partial hearing loss;^2-4^ this electro-acoustic stimulation significantly improves heaving outcomes, particularly for complex listening tasks.^5, 6^ While CIs and other neural implants provide beneficial and restorative functions, the foreign body response may impede the implant function over time and induce other deleterious effects.^7-9^ When a CI is inserted, non-specific proteins adsorb to the surface which then recruit other proteins and cells, such as macrophages.^10, 11^ One study showed that over 95% of cochleae from CI patients contained significant quantities of foreign body giant cells.^12^ In some cases, phagocytosed titanium and silicone have been found throughout the body.^13^ If cells cannot digest the foreign material, a fibrous capsule is formed around the implant. This dense tissue development around a CI can significantly impede electrical signal from reaching the target neural cells, which can dramatically reduce the hearing quality.^14-17^ Additionally, the tissue buildup and resultant scarring may even propagate fibrotic damage to more distal areas of the cochlea, resulting in partial or complete loss of residual low-frequency hearing for those with hybrid implants.^18-22^

Various solutions have been proposed to inhibit or negate the foreign body response, thus enhancing the lifetime and efficacy of medical implants. One strategy is to modify the implant surface to become more like native tissue, leading to reduction in the fibrotic response. For example, polyethylene glycol (PEG) coatings produce a more inert surface that limits binding sites for biomolecules.^23^ While PEG-derivatives are considered to be antifouling, these coatings often do not prevent deleterious long term fibrosis effects.^24, 25^ A different avenue explored for prevention of the foreign body response is the inclusion of agents such as dexamethasone, an anti-inflammatory drug, or metal nanoparticles into a coating.^26, 27^ At sufficient concentrations, these additives can mitigate the foreign body response. Unfortunately, due to the relatively short delivery timeframe, effectiveness for long-term, indwelling implants such as CIs is limited.

More recently, zwitterionic polymers have been explored as an alternative means to reduce the foreign body response.^28, 29^ In particular, zwitterionic polymer networks swell significantly while strongly binding surface water, leading to a large decrease in protein adhesion and cellular response.^30^ The positive and negative charges of the zwitterions readily interact with water, which leads to formation of a water layer that makes it difficult for biomolecules to interact and adhere to the surface. To take advantage of these antifouling characteristics, significant efforts have been devoted to modifying the surface of other materials using brush zwitterionic polymers. However, these brush polymers are quite thin, and as a linear polymer, lack durability to remain viable as a coating for high-abrasion implant situations, including CI implantation. To enhance stability, crosslinker can be added to zwitterionic systems to form a hydrogel.^31-33^ These hydrogels also may lack significant mechanical stability and thus are often not suitable coatings for metal implants or those produced from stiffer polymers. ^34, 35^ To incorporate zwitterionic hydrogels for effective antifouling and reduced fibrosis of implants such as CIs, stability of the coatings must be sufficient to withstand mechanical deformation during the coating process, throughout surgical insertion, and the implant lifetime.

Previous work has demonstrated that zwitterionic thin films can successfully be coated on biomaterials by simultaneous photografting and bulk photopolymerization, using Type II and Type I photoinitiators, respectively.^36^ Photografting allows rapid and spatially-controlled covalent attachment of zwitterionic hydrogels to the sample surface which significantly decreases biofouling.^37-39^ A compromise between mechanical integrity and antifouling capability was observed as a function of crosslink density for the photografted zwitterionic hydrogel thin film system.^31^ Additionally, CIs undergo bending and encounter hard tissue during implantation,^40^ requiring that coatings be robust and remain attached. This work focuses on the application of zwitterionic thin films to CI biomaterials to enhance antifouling and lubricity. The durability of the photografted zwitterionic hydrogel coatings was examined during exposure to extreme but relevant conditions including desiccation under ambient conditions, increasing normal forces, and bending. Polydimethylsiloxane (PDMS), typically the housing for electrode arrays of CIs, was coated with zwitterionic hydrogels, and lubricity was compared between coated and uncoated samples to determine if surface properties are maintained. Durability under bending forces and desiccation of coated PDMS samples was also examined. Material properties and integrity were also determined before and after explantation to confirm long-term durability *in vivo*. This work demonstrates that CI biomaterials can successfully be coated with zwitterionic hydrogels that remain intact and adhered during implantation and throughout the functional lifetime of the implant.

## 2. Methods

### 2.1 Monomer Solutions

Photoinitiator 2-hydroxy-1-[4-(2-hydroxyethoxy) phenyl]-2-methyl-1-propanone (HEPK, Sigma Aldrich) was dissolved into deionized water to obtain an approximately 0.077 wt% solution. The two zwitterionic monomers used (chemical structures shown in figure 1) were sulfobetaine methacrylate (SBMA, 2-methacryloyloxy)ethyl]dimethyl-(3-sulfopropyl)ammonium hydroxide, Sigma Aldrich) and carboxybetaine methacrylate (CBMA, 3-[[2-(methacryloyloxy)ethyl]-dimethylammonio]propionate, TCI Chemicals). Poly(ethylene glycol) methacrylate (PEGMA, 400 g mol^-1^, Polysciences) and 2-hydroxyethyl methacrylate (HEMA, Sigma Aldrich) were used as non-zwitterionic monomer controls. Poly(ethylene glycol) dimethacrylate (PEGDMA, 400 g mol^-1^, Polysciences) was used as the crosslinker (chemical structure shown in figure 1) for all monomer systems. Monomer and crosslinker were added at various ratios totaling 35 wt% of the prepolymer solution, with the remaining 65 wt% composed of the water/HEPK mixture. Monomer solutions and resultant hydrogel films are identified with the percent of the total monomer that is PEGDMA (crosslinker).

**Figure 1.**
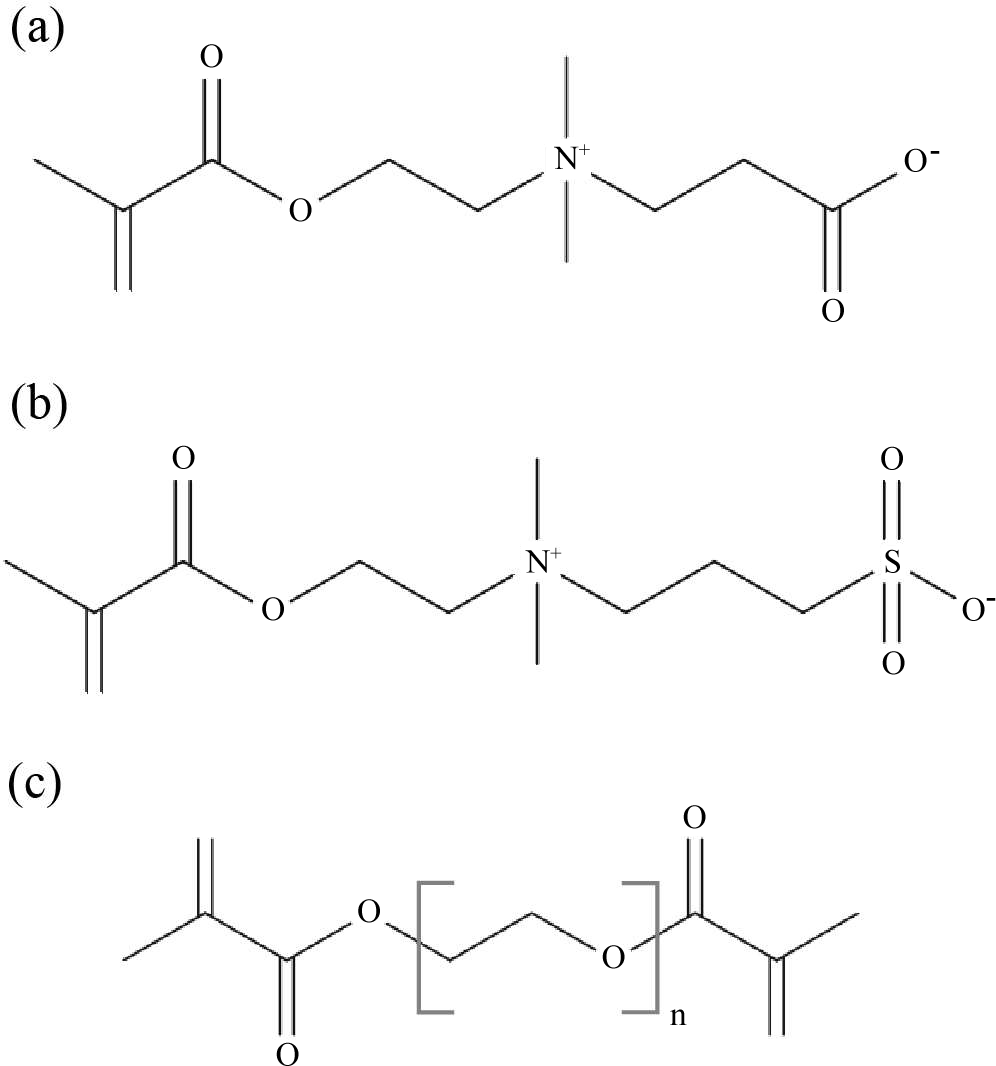
Chemical structures of (a) sulfobetaine methacrylate (SBMA) (b) carboxybetaine methacrylate (CMBA) and (c) poly(ethylene glycol) dimethacrylate (PEGDMA).

### 2.2 Coated Samples

Reinforced medical grade PDMS (Bentec Medical) was cut into disks (2.54 mm thick with 25 mm diameter or 0.016 mm thick with 12 mm diameter) or rectangles (35x75x1 mm). PDMS samples were soaked in a 50 g benzophenone (Sigma Aldrich)/L acetone solution for one hour. The PDMS samples were then removed from the solution and vacuum dried for at least 20 minutes to evaporate any residual acetone. Prepolymer solution (20 µL for 25 mm diameter disks, 5 µL for 12 mm diameter disks, 200 µL for rectangles) was pipetted onto the benzophenone-treated PDMS and dispersed with glass coverslips (Fisher Scientific). The solution was then polymerized using an Omnicure S1500 lamp at 30 mW cm^-2^ for 10 minutes under full spectrum light (300-520 nm) to simultaneously photograft and form the hydrogel coating.^36^ Coated PDMS samples were placed in Dulbecco’s phosphate buffered saline solution (PBS, Gibco, Thermo Fisher Scientific) for at least 24 hours to allow hydrogel coatings to swell to equilibrium.

### 2.3 In vivo implantation

To examine *in vivo* durability, coated and uncoated PDMS samples were implanted subcutaneously in mice for 16 days or six months and then explanted. For 16 day samples, representative images were collected using confocal microscopy, while for six month implants, coatings were imaged using scanning electron microscopy.

### 2.4 Tribometry

Lubricity of disk samples was evaluated using a pin-on-disk tribometer (TRB^3^, Anton Paar) as previously described.^41^ Typically, each sample was examined for 20 cycles (∼8 minutes) at a rotational speed of 1.3 mm s^-1^ using the tribometer liquid setting while immersed in PBS. Samples were subjected to a one N normal force using a sapphire probe. A coefficient of friction curve was generated for each sample (see supplemental figure 1(a) as an example) including the mean coefficient of friction for the experimental run time. To determine longevity of samples, coefficient of friction data were collected over 2500 cycles (∼1000 minutes) for coated and uncoated samples. The values for all coated samples were normalized to that of bare PDMS (uncoated). To examine varying force, the normal force applied for each run was varied between 1 and 15 N.

To investigate effects of rehydration, samples were initially swollen to equilibrium followed by exposure to ambient conditions for 24 hours and then under vacuum for 10 minutes to achieve complete desiccation. The samples were then rehydrated in PBS for 24 hours prior to testing. Coefficient of friction was determined using the tribometer and compared to control samples which were swollen to equilibrium without desiccation. Further, to ascertain the effects during desiccation on the coefficient of friction, coated PDMS samples, initially at equilibrium in PBS, were tested without additional liquid in the tribometer sample holder to allow water evaporation. The tribometer probe exerted normal forces while the coefficient of friction was measured for 90 minutes or until the coefficient of friction reached an asymptotic maximum value. Using the range of minimum to maximum coefficient of friction, T10 and T90 values were calculated where T10 is the time for the coefficient of friction to increase to 10% of the total range for the sample and T90 the time to reach 90% of the range (see supplementary figure 1(b) tribometry results with T10 and T90 values indicated).

#### 2.4.1 Tribometry ex vivo and after implantation

Tissues were harvested from guinea pigs following approved methods by the University of Iowa Institute of Animal Care and Use Committee (IACUC #9092245). Selected tissues were cut into 1x1 cm pieces. The explanted tissues were then used to cover a steel probe head prior to tribometry, as illustrated previously.^41^ Coefficient of friction was measured between the tissues and coated-PDMS surfaces. For an additional control, the coefficient of friction for dermis on dermis was measured after fixing dermis tissue to PDMS using tissue glue. Coefficient of friction for uncoated PDMS with each tissue was also measured. Smaller diameter (12 mm) PDMS disks, both coated and uncoated, were implanted subcutaneously in 10 week old CBA/J mice for three weeks, following protocols approved by the University of Iowa Institutional Animal Care and Use Committee. After removal from the mice, lubricity was measured with samples immersed in PBS. For comparison, pristine samples, identical to those implanted but simply stored in PBS for the duration of implantation, were also examined.

### 2.5 Mandrel bend

Flexural failure of coated rectangular samples was examined using a mandrel bending apparatus.^42, 43^ All samples not containing 100% crosslinker withstood failure for bending around all diameter (2-32 mm) cylinders when fully hydrated. To ascertain the effect of dehydration, samples were subjected to ambient conditions allowing desiccation and bent around a 5 mm diameter mandrel every five minutes. The time to failure indicates the time at which cracks in the coating were observed.

### 2.6 Insertion Force of Cochlear Implant Electrode Arrays

SBMA thin films were coated on cochlear implant electrode arrays using a previously published method.^41^ In brief, electrode arrays were pretreated for one hour in an acetone solution with 50 g/L benzophenone, as above. After removal from the solvent and drying under vacuum, the implants were inserted into rigid cylindrical sleeves of transparent PDMS (inner diameter 0.76 mm) filled with a 10% crosslinker monomer solution. To enable adequate dispersion of the hydrophilic monomer solution over the hydrophobic PDMS surface, 0.8 wt% surfactant (dimethylsiloxane-acetoxy terminated ethylene oxide block copolymer, 75% nonsiloxane, Gelest, Inc.) was included. This system was exposed to UV light at 30 mW cm^-2^ (Omnicure S1500 lamp, 320-500 nm) for five minutes in an oxygen-free environment. Following removal of the sleeve, the system was UV cured for an additional 10 minutes to ensure complete polymerization. Coated arrays were soaked in a fluorescein disodium salt solution to allow for visual comparison with uncoated arrays. Both coated and uncoated cochlear implant electrode arrays (Cochlear Slim Straight and Advanced Bionics Slim J with lengths of 23-25 mm) were inserted into human cadaveric cochleae. The cochleae were mounted on a force transducer to assess the increase in force associated with an insertion run.^41^ The raw output was reported over time for each implant.

## 3. Results

### 3.1 Durability of thin films in vivo

To determine the attachment and film integrity of photografted zwitterionic thin film hydrogels *in vivo*, PDMS samples coated with zwitterionic hydrogels of varying crosslink density were subcutaneously implanted in mice. After incubation for two weeks or six months, samples were excised and imaged. Films, independent of crosslink density, remained intact with little to no evidence of delamination via scanning electron microscopy (figure 2(b)-(e)). The films with 2% crosslinker appeared to have a more irregular surface, whereas the surface of films with 5-31% crosslinker were relatively smooth with uniform thickness. Confocal z-stack imaging of fluorescein immersed samples implanted for 16 days (figure 2(f)-(g)) also showed that thin film hydrogels incorporating 10% crosslinker retained mechanical integrity without any visible defects or deformation.

**Figure 2.**
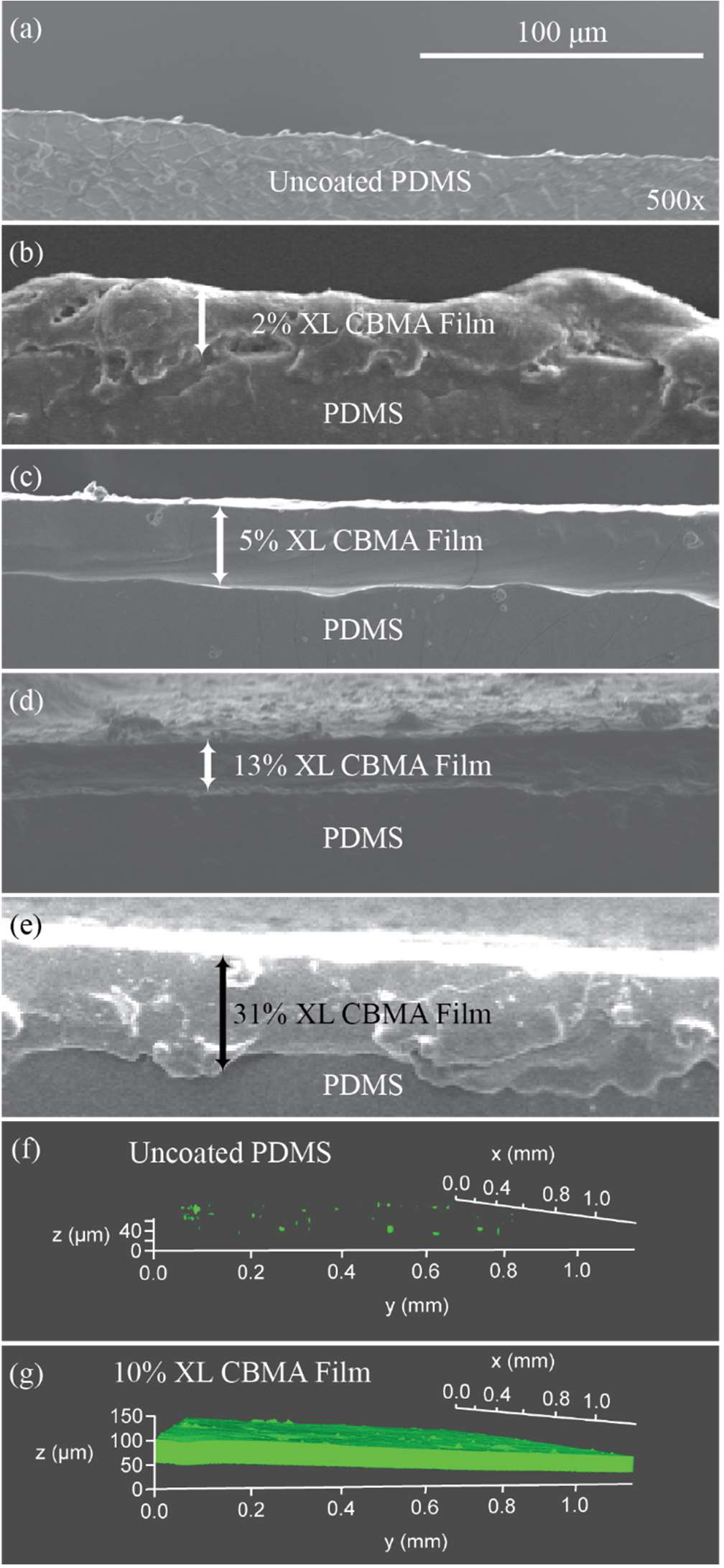
Scanning electron microscope cross-section images of six-month implants of (a) uncoated PDMS and (b)-(e) CBMA-coated PDMS with indicated crosslinker (XL) percentages. Scale bar and magnification for (a)-(e). Confocal microscopy z-stack images of 16-day implants immersed in fluorescein (green) are also shown of (f) uncoated and (g) 10% XL CBMA-coated PDMS.

### 3.2 Lubricity following implantation in mice

To determine if zwitterionic hydrogel coatings remain stable without changes in frictional resistance, coated and uncoated PDMS samples were implanted subcutaneously into mice for three weeks. The coefficient of friction was then determined for explanted samples and those simply stored in PBS for three weeks. A slight, but statistically insignificant, increase in coefficient of friction was observed from the pristine to explanted uncoated samples (figure 3).

**Figure 3.**
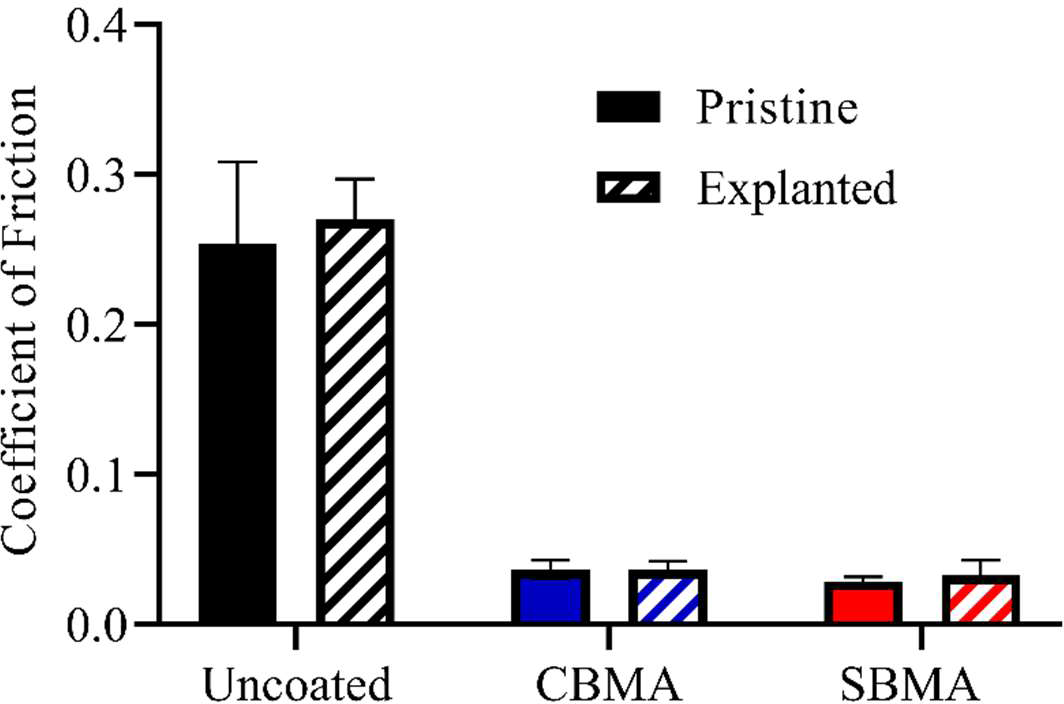
Coefficient of friction of uncoated, CBMA-coated, and SBMA-coated PDMS f or pristine samples and samples explanted from mice after three weeks incubation measured with tribometry using PBS as immersive solution and sapphire probe setup. Error bar indicates standard error of mean for n≥4.

Additionally, no difference was evident between implanted and pristine samples for either CBMA- or SBMA-coatings, all exhibiting approximately 90% reduction in coefficient of friction relative to uncoated PDMS. Taken together, these results showed that the lubricity imparted by zwitterionic hydrogel coatings was not affected by implantation and *in vivo* incubation.

### 3.3 Lubricity of coatings relative to guinea pig tissue

To determine the impact of zwitterionic hydrogels on lubricity with biological systems, the tribometer probe was covered with various freshly excised guinea pig tissues. This modification provided a direct perspective on the effect of hydrogel coatings on the lubricity between implants and surrounding tissue, especially as might be relevant during implantation. The coefficient of friction was examined between a variety of freshly excised guinea pig tissues and photografted 5% crosslinker SBMA coatings as well as uncoated PDMS controls as shown in figure 4(a). The zwitterionic coating induced at least a 90% reduction in friction relative to uncoated PDMS for all tissues. Slightly higher coefficient of friction values were observed between the zwitterionic hydrogel and dermis and trachea tissues, similar to that observed simply with the steel probe. To determine if changes in hydrogel composition affected the lubricity with tissue, the coefficient of friction between dermis tissue and various hydrogel coatings, including SBMA with different crosslink densities in addition to CBMA and other hydrogel systems with 5 wt% crosslinker, was investigated (figure 4(b)). Dermis was chosen as the representative tissue, both for ease of use and because it showed the highest coefficient of friction of the tissues tested. A slight increase in coefficient of friction was evident as the crosslink density increased for SBMA coatings, similar to trends observed in previous work for the zwitterionic hydrogel system using an uncovered sapphire probe.^31^ At 5% crosslinker, both CBMA and PEGMA coatings showed a similar reduction in coefficient of friction (∼95%) as SBMA relative to uncoated PDMS with dermis tissue. Conversely, HEMA with 5% crosslinker and PEGDMA coatings induced much higher friction, with the highly crosslinked PEGDMA coating reducing the coefficient of friction about 50% and HEMA demonstrating a similar value to uncoated PDMS. Interestingly, when dermis was tested as both probe covering and substrate, the coefficient of friction was about 30% higher than that of dermis with uncoated PDMS, showing that the zwitterionic hydrogels dramatically increase the lubricity beyond what might even occur between native tissue in the body.

**Figure 4.**
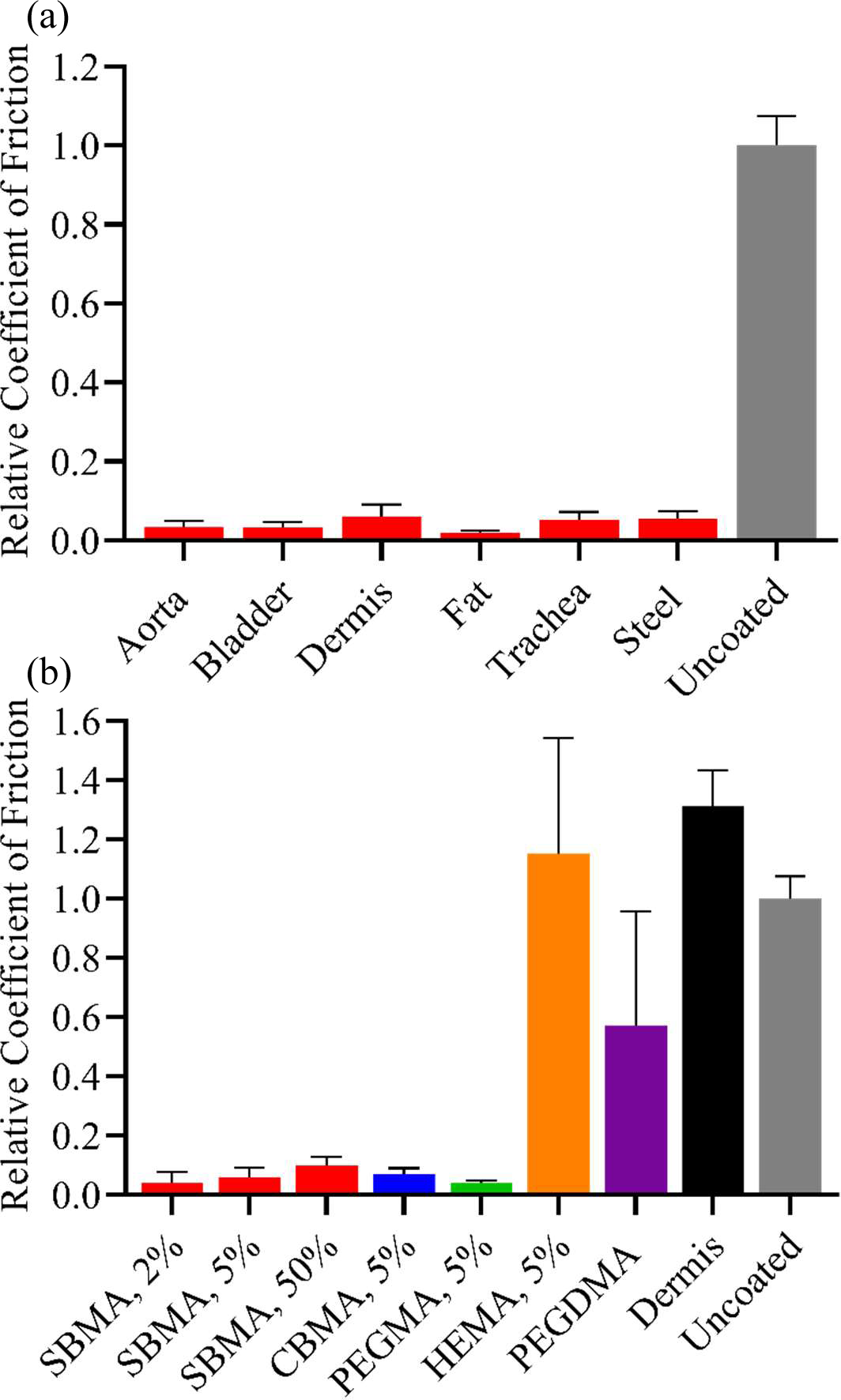
Coefficient of friction relative to uncoated PDMS for (a) 5% SBMA hydrogel coated PDMS against various guinea pig tissues or non-tissue covered steel probe and (b) different hydrogel coatings (percents are crosslinker %) or dermis itself against the dermis covered probe. Uncoated PDMS value for dermis-covered probe 0.301. Error bar indicates standard error of mean for n≥3.

### 3.4 Coating stability with desiccation

To examine the impact of desiccation, the lubricity of zwitterionic hydrogel coatings was investigated as a function of drying time and crosslinker concentration. Hydrogels were swollen to equilibrium in PBS. Frictional resistance was thereafter measured immediately after removal from solution while allowing water to evaporate in ambient conditions. To quantitatively compare when hydrogel coatings start to lose lubricity with decreased hydration, T10 and T90 (i.e., times at which the coefficient of friction has reached 10% and 90%, respectively, of the total range of observed values; see supplemental figure 1(b)) were determined. CBMA coatings with lower crosslink densities (figure 5(a)) showed T90 values over 60 minutes, whereas T90 values decreased to around 30 minutes with higher crosslink density films. T10 values, indicating when lubricity starts to measurably decrease, followed the same trend with shorter times at higher crosslink densities. The lower crosslink density films showed T10 values around 40 minutes while more crosslinked films lost measurable lubricity as early as 20 minutes. Similarly, SBMA coatings (figure 5(b)) required up to 60 minutes to reach the T10 threshold, again with decreasing values as crosslinker percent increases. T10 values were comparable between the two zwitterionic thin films with CBMA remaining intact for slightly longer times at low crosslink densities. PEGMA, a non-zwitterionic hydrogel, (figure 5(c)) exhibited T90 times up to approximately 60 minutes at lower crosslink densities. While the trend was not as consistent, PEGMA T10 values decreased with increasing crosslinker percent on the same order as the zwitterionic systems. While T10 and T90 values for PEGMA were higher than CBMA and SBMA for some crosslinker percents, the actual coefficient of friction of PEGMA-coated samples was also higher initially. For example, the initial coefficient of friction for PEGMA with 2% crosslinker was 0.058 compared to 0.041 and 0.047 for CBMA and SBMA, respectively, with the same amount of crosslinker. Thus, zwitterions demonstrate capability to retain water and decrease lubricity for similar lengths of time to the PEG systems while imparting a greater reduction in overall coefficient of friction.

**Figure 5.**
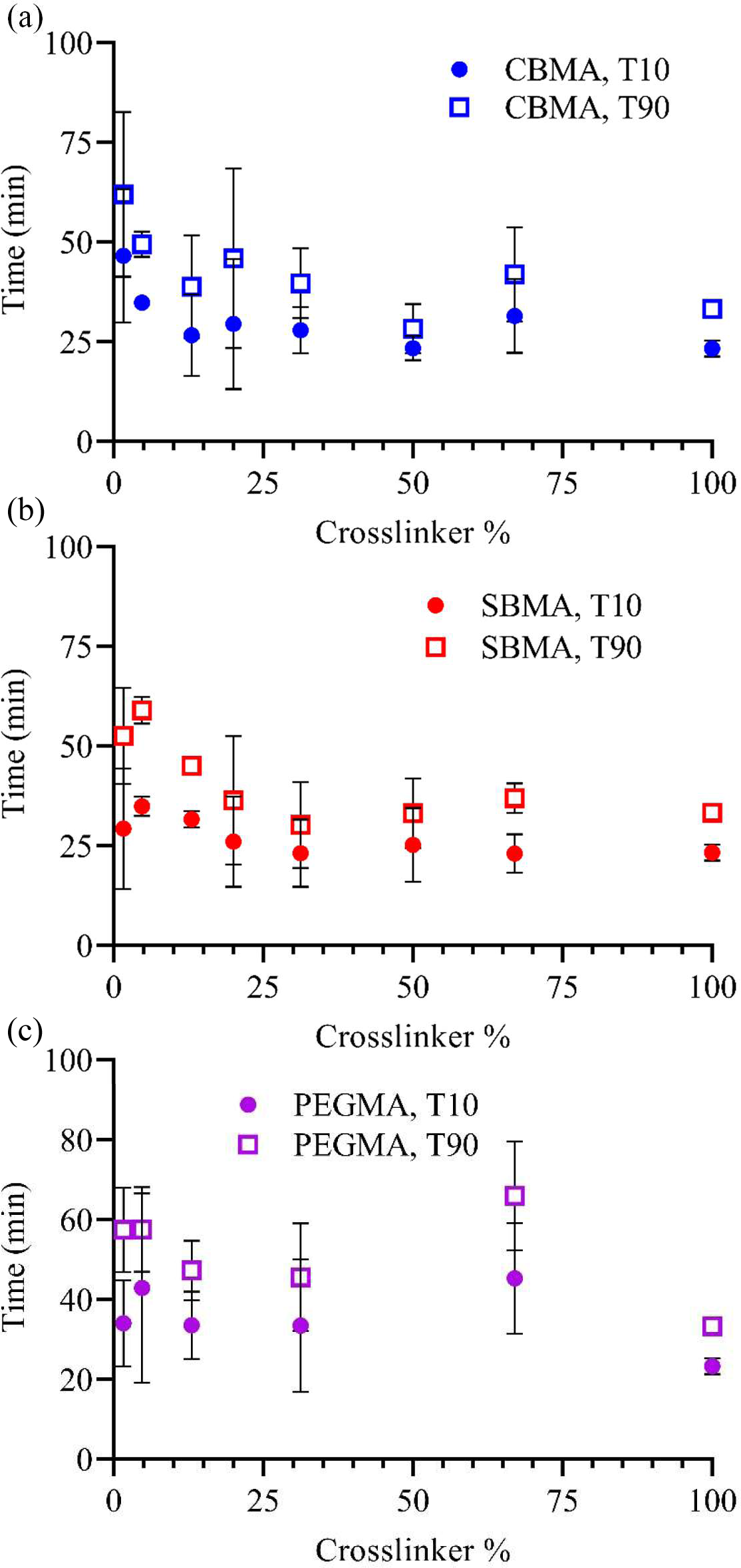
The time for (a) CBMA, (b) SBMA, and (c) PEGMA coatings to reach 10% (T10) and 90% (T90) of maximum coefficient of friction value over a range of crosslink densities. Hydrogels were swollen to equilibrium in PBS prior to testing but measured without additional PBS. Error bar indicates standard deviation for n≥3.

Hydrogel coatings were also examined for robustness when subjected to bending forces using a modified mandrel bend test. For all hydrogel coatings except 100% crosslinker (PEGDMA), completely hydrated coatings remained unaffected after bending over the complete range of diameters. To quantify the effects of bending and desiccation, coatings were exposed to ambient conditions to allow water loss. Each system was then bent around a 5-mm cylinder every five minutes until cracking was observed. Both CBMA and SBMA coatings displayed viability for reasonably long periods of time, especially at lower crosslink densities (figure 6). With five percent crosslinker, CBMA and SBMA did not exhibit signs of failure until 75 and 50 minutes, respectively. The time to failure for CBMA decreased with crosslinker percent, but coatings remained viable for up to 30 minutes even with more than 60% crosslinker. In contrast to the results discussed previously, PEGMA systems behave much differently than the zwitterionic systems when exposed to bending forces under desiccation. The PEGMA samples failed at least 25 minutes earlier than SBMA and 50 minutes earlier than CBMA at low to intermediate crosslinker percents (figure 6). The neat crosslinker (PEGDMA) as a hydrogel failed upon bending when still in the equilibrium hydration state.

**Figure 6.**
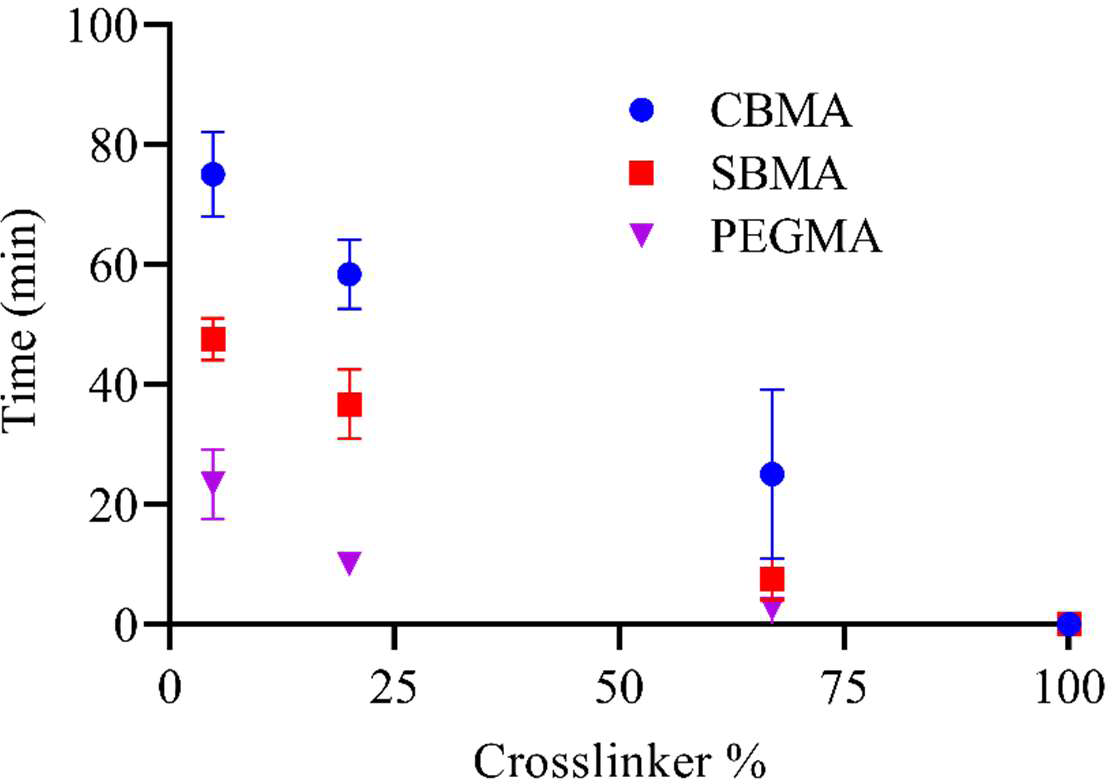
Time until failure was first noted for hydrogel coatings when subjected to 5mm diameter mandrel bend test for a range of crosslink densities. Hydrogel coatings were swollen to equilibrium and then exposed to air (allowed to dry) starting at time 0. Error bar indicates standard error of mean for n≥3.

### 3.5 Lubricity before and after desiccation

To assess the effects of complete desiccation and subsequent rehydration, the coefficient of friction for SBMA hydrogel coatings with different amounts of crosslinker before and after complete desiccation was investigated (figure 7). At low crosslink densities, the coefficient of friction was nearly identical for both conditions. As the crosslinker percent increases, the disparity between the rehydrated sample and the pristine sample begins to increase. At higher concentrations of crosslinker, the rehydrated samples exhibit coefficient of friction values similar to that of uncoated PDMS. Qualitatively, these higher crosslinked coatings were quite fragile when dried with almost complete failure and little coating remaining when the sample was rehydrated in PBS, suggesting that the rehydrated sample was primarily uncoated PDMS. While the 100% crosslinker sample did exhibit a slight increase in the coefficient of friction following drying, the difference is not as large as with the highly crosslinked zwitterionic coatings. Samples without crosslinker displayed a much higher coefficient of friction after initial swelling relative to other SBMA coatings with no significant change noted after drying and rehydration.

**Figure 7.**
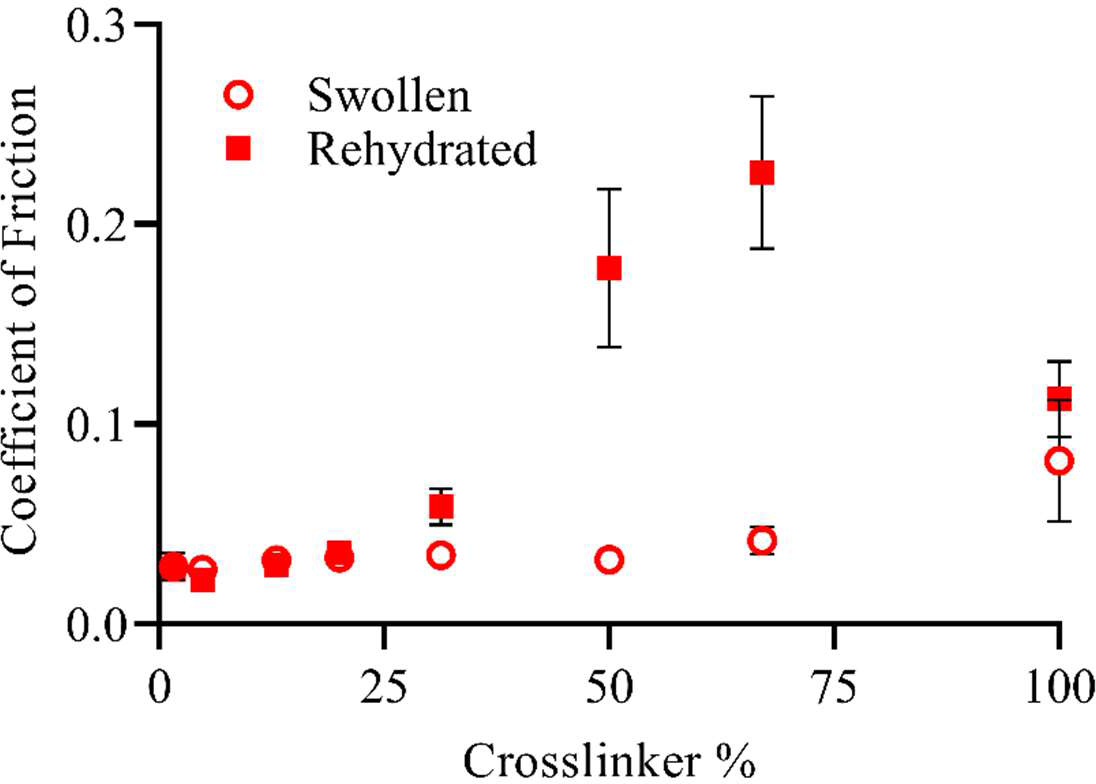
Coefficient of friction for SBMA hydrogels as function of crosslinker percent in swollen state immediately after polymerization and following complete desiccation and rehydration. The value for uncoated PDMS Error bar indicates standard error of mean for n≥4. Uncoated PDMS had an average value of 0.179.

### 3.6 Long-term lubricity

To ascertain the effect of implant duration on zwitterionic hydrogel coatings, the coefficient of friction as a function of crosslink density was measured for SBMA-coated PDMS using tribometry over 2500 cycles equating to nearly 17 hours under force. The average coefficient of friction for the first and last hundred cycles, corresponding to approximately 40 minutes of measurement, was compared to show the difference in surface properties under constant load (figure 8). For all crosslinked zwitterionic thin films, the relative coefficient of friction was not statistically different between the first and last 100 cycles. A slight, though statistically insignificant, increase in the coefficient of friction was observed for 100% PEGDMA samples from repeated loading. On the other hand, the coefficient of friction value for SBMA samples without crosslinker decreased significantly (p<0.0001).

**Figure 8.**
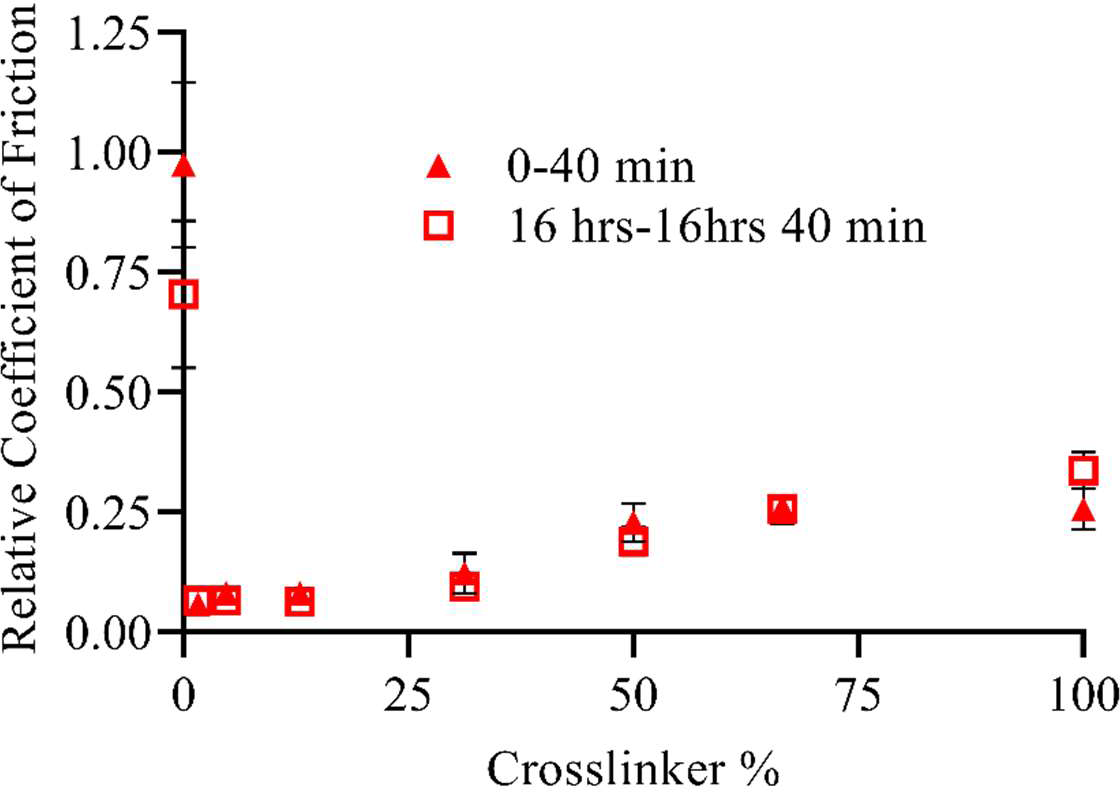
The average coefficient of friction (relative to uncoated PDMS) for the first and last 100 cycles of a 2500 cycles test of SBMAcoated PDMS as a function of crosslinker percent. The cycle time equates to a total testing period of 1000 minutes, with sampling occurring over the first and last 40 minutes. Error bar indicates standard error of mean for n≥4. Average coefficient of friction of for uncoated PDMS was 0.197.

### 3.7 Coefficient of friction with different applied normal force and speed

To approximate the range of forces that such hydrogel coatings might encounter in a biological environment, the coefficient of friction was measured under increasing normal forces for SBMA hydrogels with 5% crosslinker. As shown in figure 9, a nearly linear increase in the coefficient of friction was observed up to 10 N, raising the coefficient of friction 250%. Even with greater normal force, the coatings still maintained a coefficient of friction much lower than that of uncoated PDMS. Qualitatively, the hydrogels showed few, if any, noticeable defects at low normal force. At higher forces, however, the wear track became visible after tribometry was completed. Additionally, the lubricity of SBMA coatings was compared for probe speeds of 1 mm s^-1^ and 6 mm s^-1^. For all crosslink densities, no significant difference was observed in coefficient of friction values with increased speed (supplemental figure 2).

**Figure 9.**
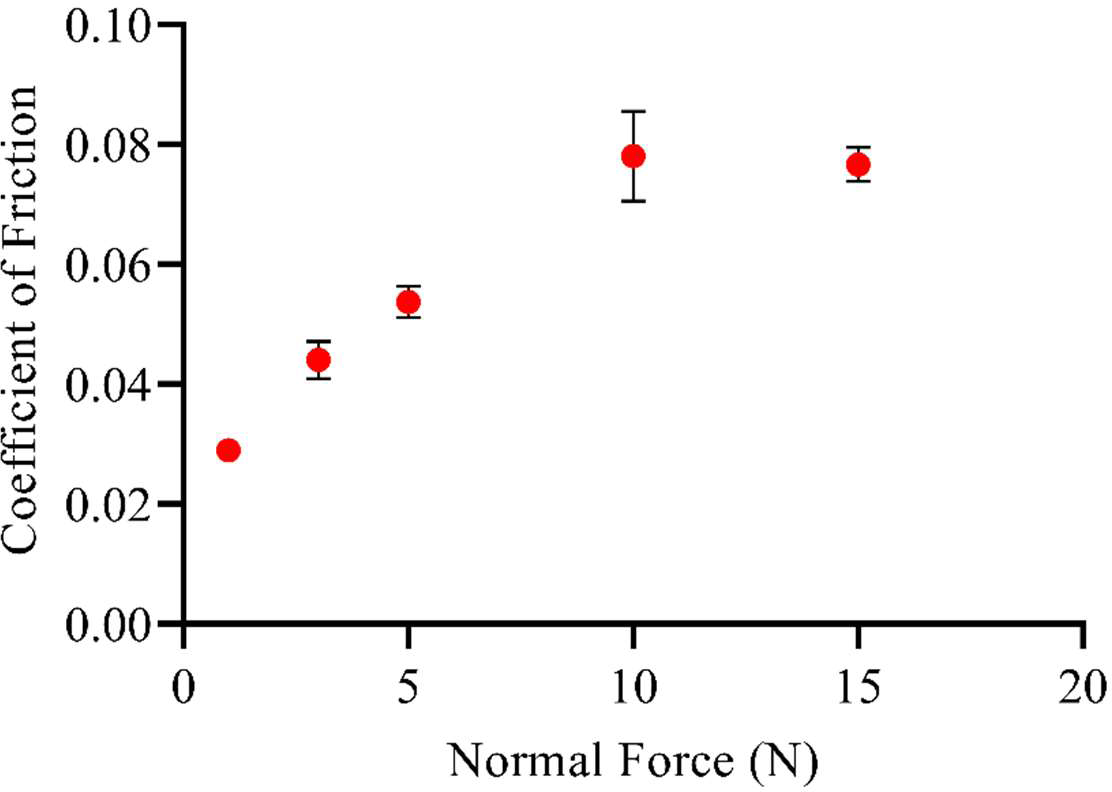
Coefficient of friction for SBMA hydrogels (5% crosslinker) with increasing normal force applied. Uncoated PDMS value for normal force 1N 0.19. Error bar indicates standard error of mean for n≥4.

### 3.8 CI Insertion force with zwitterionic coatings

Zwitterionic hydrogels were effectively coated on CI electrode arrays using simultaneous photografting and photopolymerization, as described earlier. As evident by visualization using a fluorescein dye, the hydrogel coats the CI uniformly (figure 10). To determine if these zwitterionic hydrogel coatings impact insertional forces in biological tissues, the force required to implant both coated and uncoated CI lateral wall electrode arrays into cadaveric human cochleae was investigated. Figure 11 shows representative insertion force profiles as a function of insertion time for both SBMA-coated and uncoated electrode arrays from two different manufacturers. Videos showing insertion of both coated and uncoated can be found in supplemental information. For array type I, the measured force for the uncoated array began to rapidly increase about halfway through the insertion with a maximum value of around 90 mN. On the other hand, the coated array type I experienced a gradual increase to a maximum force of only around 25 mN (figure 11(a)). The type II uncoated array required increased force from the onset with a maximum around 45 mN, while the coated array reduced the required force to approximately 35 mN (figure 11(b)).

**Figure 10.**
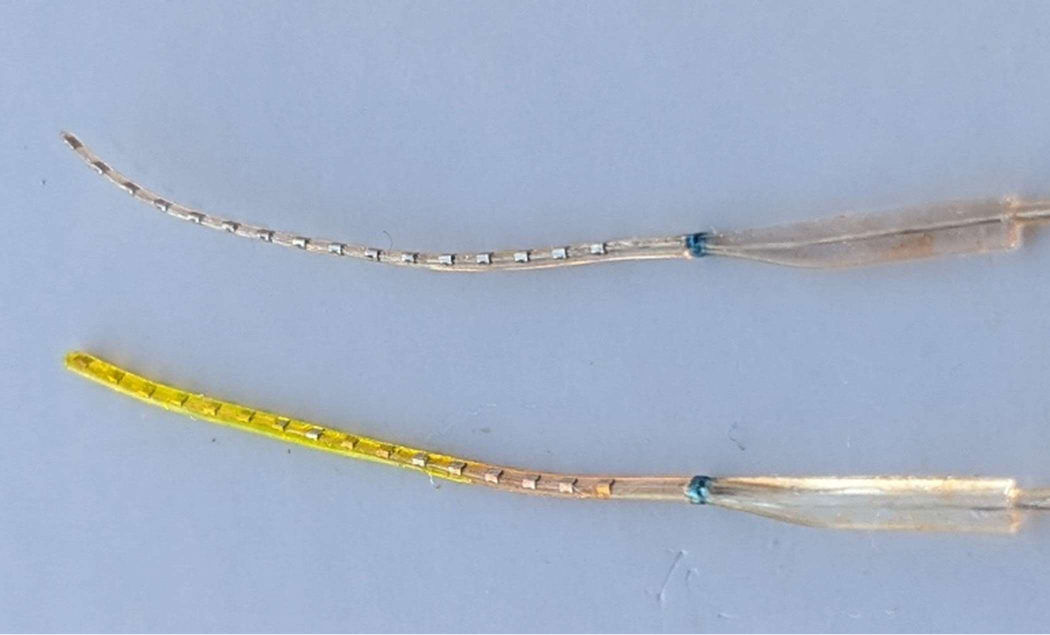
Representative image of uncoated (top) and coated (bottom) cochlear implant arrays. The coated array was soaked in fluorescein solution for better visualization.

**Figure 11.**
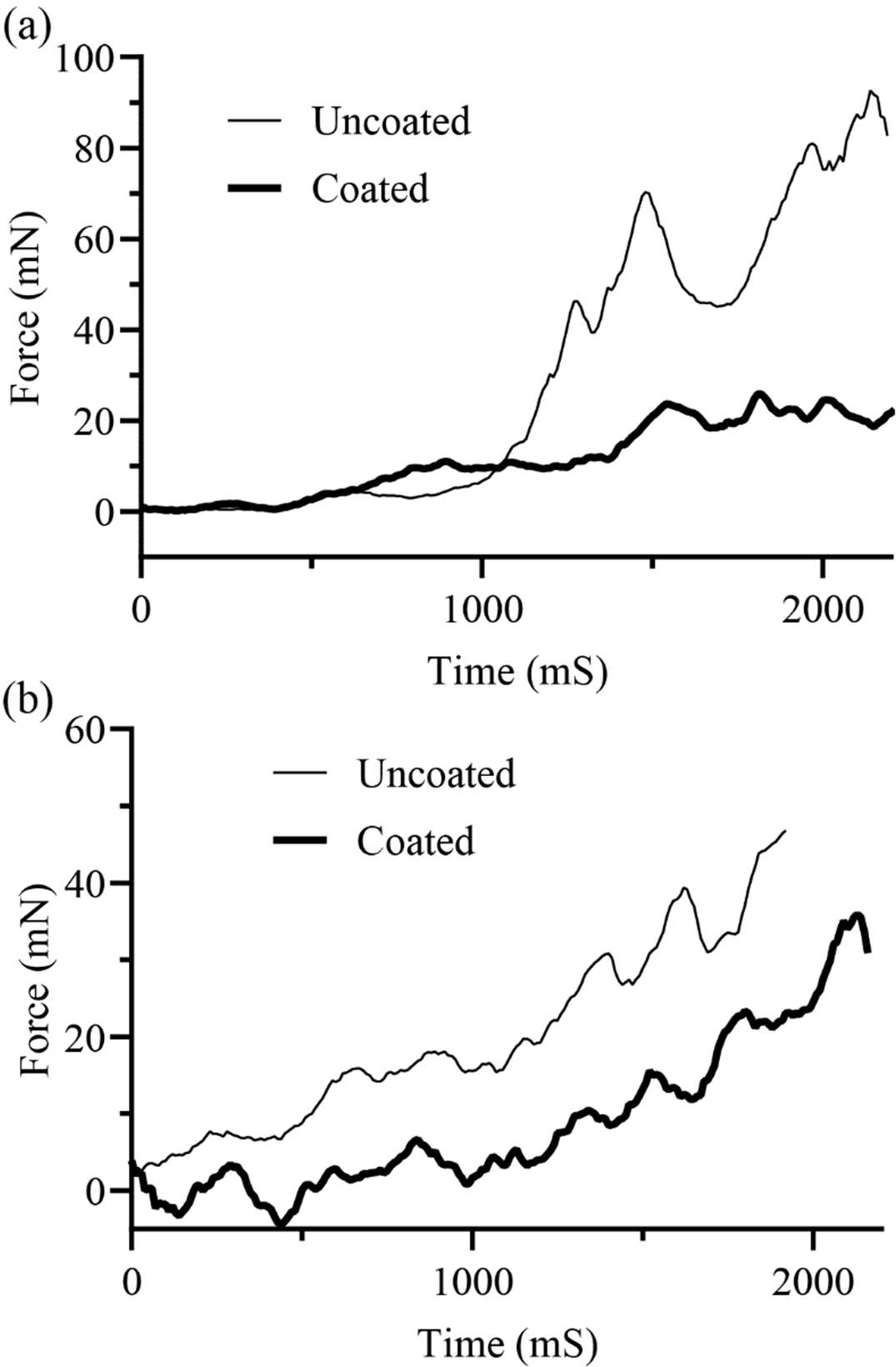
Representative force insertion profiles for uncoated (gray) and SBMA-coated (black) electrodes depicted over the time course of electrode insertion, wherein can be seen the lower maximum force and lower overall force over time for two manufacturers: (a) Array Type 1 and (b) Array Type 2.

The maximum force for each implantation was also determined as shown in figure 12. For both array types, a significant reduction (∼30%) in the maximum insertion force was determined between the uncoated and coated samples (figure 12(a)). The work of insertion was calculated using the area under the force curve for both array types, comparing uncoated and SBMA-coated implants. A decrease in the work was also observed, although the values did not statistically reflect a significant difference for coated and uncoated arrays due to high variability, especially from the uncoated systems (figure 12(b)). Interestingly, the overall deviation in both work and maximum force is substantially lower for coated versus uncoated arrays. These data support the qualitative observation that coated CI electrode arrays were inserted more easily and with less force compared to uncoated CI electrode arrays.

**Figure 12.**
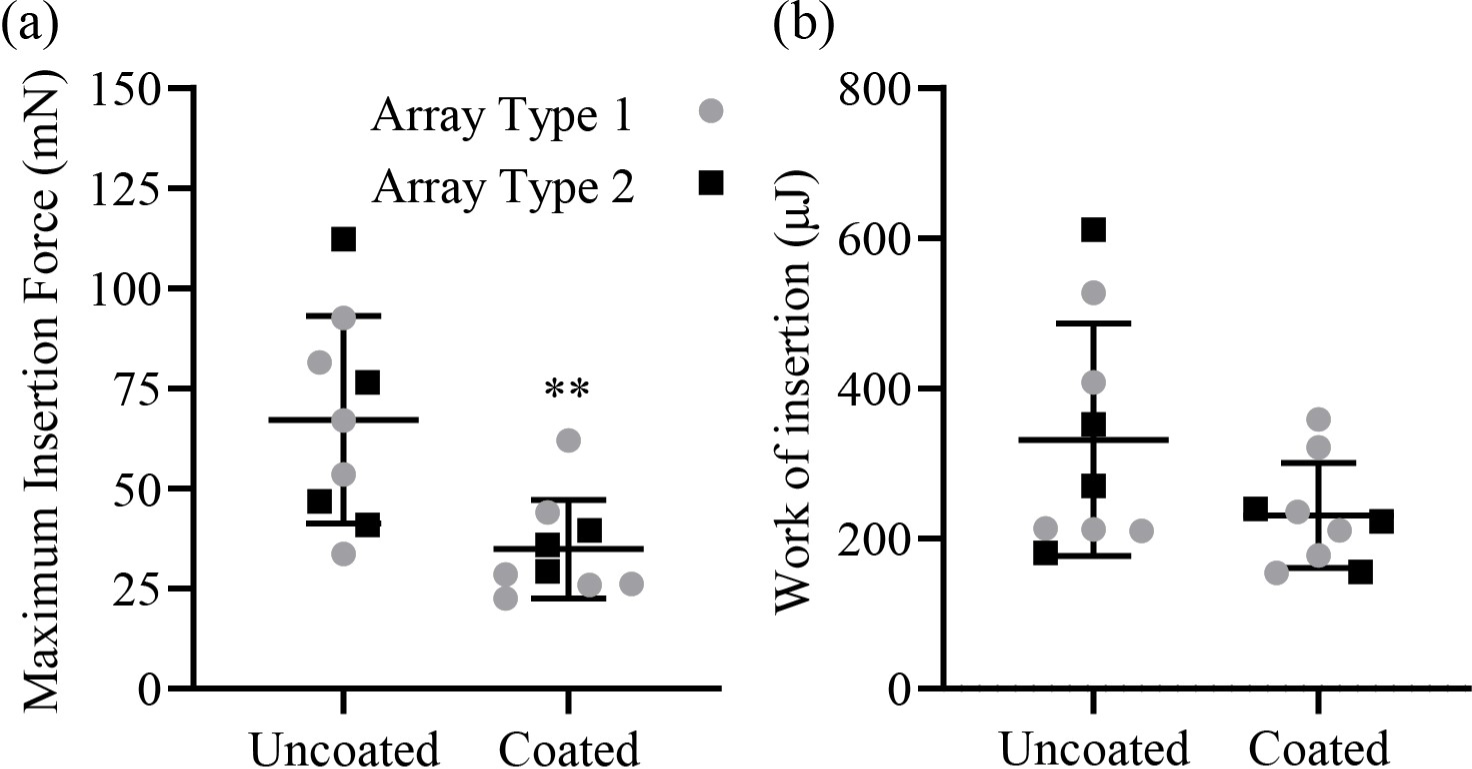
(a) The maximum force of insertion during insertions of uncoated (n = 9) and coated (n = 9) human electrode arrays. (b) Area under the curve analysis of the force over the average distance of insertion was found for insertion and averaged as a correlative measure of total work of insertion depicted in microjoules (μJ). The maximum force was significantly reduced (** p < 0.003) and overall work of insertion tended to be reduced, though this was not significant (p = 0.19).

## 4. Discussion

Hydrogels have been used extensively for a variety of biomedical applications,^44-48^ including as materials to mitigate the foreign body response.^49-53^ One major drawback of a typical hydrogel is its inherent lack of durability.^54, 55^ For a hydrogel to be reliably used for *in vivo* applications, but especially for coatings on indwelling implant systems, sufficient mechanical long-term stability is necessary to ensure viability during implantation and for the lifetime of the coated implant.^56^ Zwitterionic hydrogels photografted as biomaterial coatings have shown great promise in inhibiting the foreign body response by creation of a stable water layer leading to greater longevity and efficacy of materials and implants. To apply the unique characteristics of zwitterionic systems to various biomaterials and *in vivo* devices, the hydrogels must be sufficiently durable and maintain desirable properties throughout handling, implantation, and device lifetime. For example, CIs could greatly benefit from the antifouling capabilities of zwitterionic coatings to preserve the function of the implant and avoid loss of residual hearing from scar tissue formation. At the same time, coatings must withstand the handling prior to insertion and the normal and bending forces during insertion to be viable in this application.

To enable the use of zwitterionic hydrogel coatings, further understanding and characterization of mechanical properties and overall stability are required. Previous work has shown that, by using simultaneous photografting and photopolymerization, zwitterionic hydrogel network systems were reliably attached to PDMS surfaces.^36^ The housing of the electrode array for most CIs is composed of PDMS, a common biomaterial, which provides an excellent case study for zwitterionic hydrogel coatings biomaterials. Herein, properties including bending resistance, coefficient of friction, and *in vitro* forces were determined to understand the mechanical durability under various conditions including implantation, desiccation, increased normal force, and bending of the photografted zwitterionic hydrogel coatings. As shown by the micrographs in figure 2, zwitterionic hydrogel coatings remained intact and attached following implantation in mice for up to six months. In particular, the coatings with crosslinker amounts above 5% appeared to remain largely unchanged, indicating high-level durability for handling, surgical insertion, incubation, and removal.

Zwitterionic hydrogels are primarily of interest as biomaterial coatings to prevent biofouling due to their ability to bind water with the charged ion groups while being net neutral.^28, 57^ Another significant benefit imparted by hydrogel coatings, particularly zwitterionic hydrogels, is increased lubricity of the material surface as demonstrated by low coefficients of friction in aqueous environments.^58-60^ The significant increase in lubricity for coated surfaces should lead to less scarring and trauma during implantation, but only if the coating remains stable and retains these lubricious properties. Previous work has shown that photografted zwitterionic coatings result in an approximately 90% reduction in the coefficient of friction relative to uncoated PDMS.^31, 41, 61^ While antifouling is largely due to chemical interactions and should remain as long as the coating is still present, lubricity is more dependent on the mechanical integrity of the film. Additionally, the lubricity is reflective of the hydration state and indicative of the long-term stability of the coating. Thus, coefficient of friction was examined under various settings to determine both the inherent effects of different conditions on lubricity, as well as the implications for hydration and further stability.

The long-term stability of the coatings was further evaluated by examining lubricity of samples that had been explanted after implantation (figure 3). Both CBMA and SBMA coatings experienced no significant change in coefficient of friction after implantation for three weeks, both showing a large and consistent reduction relative to PDMS. No significant difference was observed between pristine and explanted samples, demonstrating that the zwitterionic hydrogels were stable and maintained integrity through implantation and explantation. Therefore, the large reduction in frictional resistance observed for biomaterials coated with zwitterions should remain following implantation and long-term use. As lubricity is directly related to the hydration state, these results also suggest that the zwitterionic hydrogels retain similar levels of hydration following implantation. Hydration has been directly related to the antifouling capacity indicating that these materials should also have minimal interaction with biological moieties as shown elsewhere.^62^

While enhanced lubricity could lead to less scarring while in the body and serves as a useful marker for other hydration properties, the innate decrease in friction should also lead to less trauma during insertion. For many implants, the trauma from insertion may cause deleterious effects as much as, if not more than, the innate immune response from the presence of a foreign body. For example, CI implantation led to loss of residual hearing and neo-ossification in conjunction with evidence of insertion trauma for about 50% of patients over two case studies.^63, 64^ One study found that hearing outcomes were improved when insertional trauma was minimized,^63^ suggesting that decreasing insertion trauma with increased lubricity alone will provide great benefit.

For coefficient of friction measurements, a steel or ceramic probe is typically moved across a surface of interest and the frictional force resisting movement is measured. Using such probes thereby results in measuring lubricity relative to steel or ceramic, not biologically relevant tissues. To understand if the innate lubricity of the hydrogels translates well to biological systems, the coefficient of friction between various tissues and zwitterionic coatings was examined. Such a comparison would be particularly valuable to assess the impact of coated hydrogels, especially given the resistance to insertion and the associated trauma for CIs. Interestingly, for the various guinea pig tissues tested, the coefficient of friction was dramatically reduced relative to uncoated PDMS for SBMA coatings at low crosslinker percentages (figure 4). Even when the crosslinker percentage was increased, significant reductions in friction were still observed. The coefficient of friction for dermis tissue on dermis was much higher than for the zwitterionic coatings on dermis, suggesting that the zwitterionic coatings reduces frictional resistance relative to both biomaterials and native tissue. These results also indicate that zwitterionic hydrogel coatings should reduce the friction experienced during implantation for CIs, thus significantly decreasing the trauma experienced by surrounding tissues.

All implants and implant materials are exposed to various forces, whether during insertion or within the body. Materials used for indwelling implants are sufficiently durable to withstand minute changes within the body so basic properties and performance are not affected. Because hydrogels are characterized by high water content, properties will largely be determined by the degree of hydration and surrounding environment. Therefore, loss of water will lead to significant changes in properties, such as lubricity, which could interfere with effective implantation and function. Hydrogels swell less with increasing crosslink density, which leads to an increase in some mechanical properties but is accompanied by decreased flexibility.^31, 65, 66^

To ensure that the hydrogels remained sufficiently hydrated and durable during handling, mechanical stability, as indicated by lubricity, was examined during desiccation as shown in figure 5. The T10 value was indicative of when the coating first began to dry/fail, whereas T90 occurred when the coating no longer imparted significant added lubricity to the coated PDMS. Both values indicate stability of the film and give benchmarks of how long a coating remains viable while exposed to air. T10 provides information regarding the time a film could be exposed to ambient conditions without significant loss of lubricity, e.g., how long a coated CI could be removed from solution prior to implantation without significant loss of coating surface properties. T90, on the other hand, indicates when the coating becomes largely dehydrated and represents the maximum time a coating could be exposed to air to maintain any additional lubricity.

As expected, lower crosslink density hydrogel coatings resisted failure longer than those with higher crosslink density. The films proved to be quite durable with lubricity remaining relatively constant for extended periods of time. For lower crosslink density zwitterionic films, which also have greater resistance to fibroblast and macrophage adhesion,^31^ lubricity was maintained for up to 60 minutes, which is significantly longer than would typically be required for handling and implantation. Both zwitterionic monomer systems within the range of crosslink percentages examined did not show significant signs of drying (T10) until at least 20 minutes. CBMA hydrogels at low crosslink density especially showed marked resistance to initial desiccation (T10), likely due to the greater propensity of CBMA to bind water molecules.^30^ The T10 and T90 values for both SBMA and CBMA were comparable to those measured for PEGMA hydrogels, suggesting that the zwitterionic films are durable and remain hydrated in similar fashion to well-studied PEG hydrogels. Thus, surface properties of zwitterionic hydrogels would be maintained for up to 40 minutes with ambient exposure.

Many biomaterials bend during implantation or while within the body. Coatings must remain attached and withstand both bending and normal forces to maintain zwitterionic functionality. CI electrode arrays, in particular, undergo significant bending when inserted to conform to the coiled cochlear structure, so withstanding such forces is critical for successful materials. Mandrel bend results (figure 6) confirmed the stability of the coatings in a hydrated state while also determining the time these zwitterionic hydrogel coatings remain hydrated and durable during bending even when exposed to air. Because all zwitterionic hydrogels examined remained intact with bending when swollen, insertion should not cause delamination or cracking, even at extreme angles. The CBMA hydrogels at low crosslink density lasted for well over 70 minutes in air before failure, whereas SBMA films were still viable for at least 50 minutes. The zwitterionic coatings showed greater water binding and hydration in comparison to PEGMA films which failed due to desiccation much faster. Sufficient hydration is key in allowing zwitterionic hydrogels to remain viable under bending forces, providing another example of the advantages for zwitterion coatings over PEG systems which retain less water over time.

The effects of complete desiccation on the coefficient of friction were also investigated. Through handling before implantation, hydrogel coatings might become desiccated. To determine the level of recovery from such situations, the integrity of the film and lubricity were analyzed after drying and rehydration with different crosslinker concentrations. Samples were completely desiccated before rehydrating to the equilibrium state, whereas typically coatings are hydrated to equilibrium directly following polymerization. For coatings containing less than 25% crosslinker, the coefficient of friction was basically identical between the desiccated/rehydrated and pristine samples, demonstrating that lower crosslink density zwitterionic coatings are sufficiently stable to undergo desiccation without loss of lubricity and integrity, as shown in figure 7. As the crosslinker percent increased, flaking of the coating was observed during the desiccation period, which was reflected in greatly increased frictional resistance following rehydration. Even though the zwitterionic coatings swelled much more at lower crosslinker percents,^31^ desiccation and rehydration did not damage the hydrogel network. On the other hand, for intermediate to high crosslink densities, the coatings were brittle and did not exhibit sufficient integrity during rehydration. Although the PEGDMA (100% crosslinker) film saw an increase in coefficient of friction, the difference was not as great as the highly crosslinked zwitterionic films. This difference can be attributed to lower initial swelling due to the absence of zwitterionic moieties, so complete drying does not induce as much of a physical change. Thus, at lower crosslink densities, the zwitterionic network is much more flexible, even with water removal and rehydration. These results provide a range of crosslinker percents for zwitterionic coatings that are sufficiently stable to maintain film integrity and high lubricity even if desiccation does occur at some point following the coating process. While more loosely crosslinked films do show the ability to rehydrate and maintain lubricity if not stressed in other ways, desiccated coatings did fail if exposed to bending or normal forces.

While desiccation will affect utility before implantation, stability of the coating must also be maintained for the lifetime of the implant. Some implants may be intended for short-term use, but many implants, including CIs, are intended to be permanent. For zwitterionic coatings to be viable material components, they must match the longevity of the coated implant. To assess the long-term stability, forces were applied for extended periods of time while monitoring the coefficient of friction. Figure 8 demonstrates that the coefficient of friction relative to PDMS remains stable over extended time frames for SBMA coatings across the range of crosslink densities. Some degree of crosslinking is necessary to withstand the force applied during tribometry, as without crosslinking the grafted polymer did not significantly increase lubricity compared with an uncoated PDMS surface. Although this brush-like coating did appear to become somewhat more lubricious overtime, this anomaly is likely due to the uncrosslinked coating failing and covering the probe tip leading to a slight decrease in the coefficient of friction. Conversely, even with higher amounts of crosslinker, the initial and final coefficient of friction values of zwitterionic coatings were not significantly different showing high degrees of durability for the prolonged time.

Additionally, the effect of probe speed was investigated to determine if insertion rate would impact stability. No significant difference was noted in the measured coefficient of friction with increasing speed from 1 to 6 mm/s of the applied force from the probe tip. For most implants, the greatest movement will be experienced during insertion. Kontorinis et al. reported that as insertion rate was increased from 0.17 mm/s to 3.3 mm/s for standard CIs, both mean and maximum force also increased following a roughly linear trend.^67^ Because the coefficient of friction appears to remain approximately the same even at higher probe speeds, the zwitterionic hydrogel may negate the force differences due to insertion rate, further reducing the trauma experienced.

A third factor, along with duration and speed, which may impact durability of the hydrogel coating is force magnitude. Variable forces will be encountered during implantation and while in the body. While some biomaterials are intended for short-term use, such as catheters, many are intended to be permanent. Thus, the implant life can vary widely, as well as how the implant interacts with the body once implanted. Implants may go through repeated interaction with harder surfaces, such as bone, and must be sufficiently durable to withstand such challenges for the life of the implant. An increase of force will typically increase the coefficient of friction for hydrogel coatings. At critical forces, the coating will fail and delaminate from the surface as indicated by the coefficient of friction approaching that of the uncoated substrate. The effect of increasing force was quantified for SBMA coating with 5% crosslinker, a composition that has demonstrated an appropriate balance between biological efficacy and mechanical properties.^31^

Previous research has shown the linear relationship between normal force and frictional force, particularly for skin tissue.^68^ Thus as expected, the increase in normal force applied to the zwitterionic hydrogel coatings led to greater coefficient of friction values (figure 9). Even with a 15 N normal force, the coefficient of friction values remained well below those of uncoated PDMS. While PDMS did not experience as significant of an increase in coefficient of friction over the range of forces, the coefficient of friction for the zwitterionic hydrogels was consistently less than half that of PDMS for the entire force range. Notably, at higher forces, wear tracks developed in the hydrogel coatings indicating some loss of integrity. However, these results should not affect CI usage, since maximum forces during CI implantation ranging between 0.18 and 0.42 N.^67^ The majority of implants, similarly, should not experience forces anywhere near this magnitude. On the other hand, the effect of increasing force should be considered, especially if used in load bearing implants.

As demonstrated, zwitterionic hydrogels impart increased lubricity and exhibit significant durability under a variety of conditions. During CI implantation, insertional and frictional forces from interactions with the PDMS housing of the CI are transferred to the surrounding tissue,^40, 69^ leading to trauma and scarring. With the decreased frictional resistance from the coatings, it is reasonable to believe that insertional forces should also decrease with zwitterionic coatings. To determine the impact of these systems during implantation, the force was measured to insert zwitterion coated CI electrode arrays into cadaveric cochleae which should be indicative of the forces experienced by the surrounding tissue during implantation. Zwitterionic coatings on electrode arrays significantly decreased the maximum force and work during insertion (figures 10 and 11). Array type 2 coated systems experienced about 20 mN less force consistently over the entire insertion time than the uncoated implant (figure 10(b)). On the other hand, array type 1 coated systems showed no significant change in force for about the first half of insertion but maintained a low force relative to an approximately 40 mN sharp increase for the uncoated array during the second half of the implantation (figure 10(a)). Thus, the cochleae experienced at least 50% reduction in the maximum insertion force and 30% reduction in the work of insertion for CBMA-coated implants relative to uncoated systems (figure 11). The difference in insertion force was also observed qualitatively, as can be seen in the videos included in supplementary information. Less force and manipulation were required with the coated arrays which should result in less trauma to the surrounding tissue. This decrease in insertional force would likely lead to substantially reduced scarring and trauma during implantation and thereby improved outcomes. Less dense or minimal scar tissue surrounding the implant may allow significantly improved signal transduction for CI systems leading to higher quality long-term hearing.

## 5. Conclusions

To successfully reduce the foreign body response to CIs or other biomedical implants, photografted zwitterionic hydrogel coatings must remain intact and viable during implantation and for the implant lifetime. This work demonstrates that zwitterionic hydrogel coatings on PDMS are sufficiently stable to withstand desiccation and both bending and normal forces. The coatings stay attached and intact following implantation. Zwitterionic hydrogels remain hydrated, flexible, and durable for up to 80 minutes under both normal and bending forces when allowed to desiccate in ambient conditions. The frictional resistance between zwitterionic coatings and biological tissue is up to 20 times lower than with uncoated PDMS, even after implantation in mice. Insertion of cochlear implant electrode arrays into cochleae showed that friction forces can be dramatically reduced when coated by zwitterionic hydrogels. These results clearly show that photografted zwitterionic hydrogel coatings on PDMS are sufficiently durable to be practically used for an array of biomedical implants and devices, including CIs, leading to potential reduction in both trauma during implantation and long-term foreign body response.

## Supporting information

Updated Supplementary Figures

Original Supplemental

## Acknowledgements

This work was supported by funding from the National Institutes of Health (grants R01DC012578, T32DC000040, and F30DC019274) and the American Neurotology Society 2020 Research Award.

