## Supplementary material for "Photografted zwitterionic hydrogel coating durability for reduced foreign body response to cochlear implants": Updated Supplementary Figures

### Supplementary Information

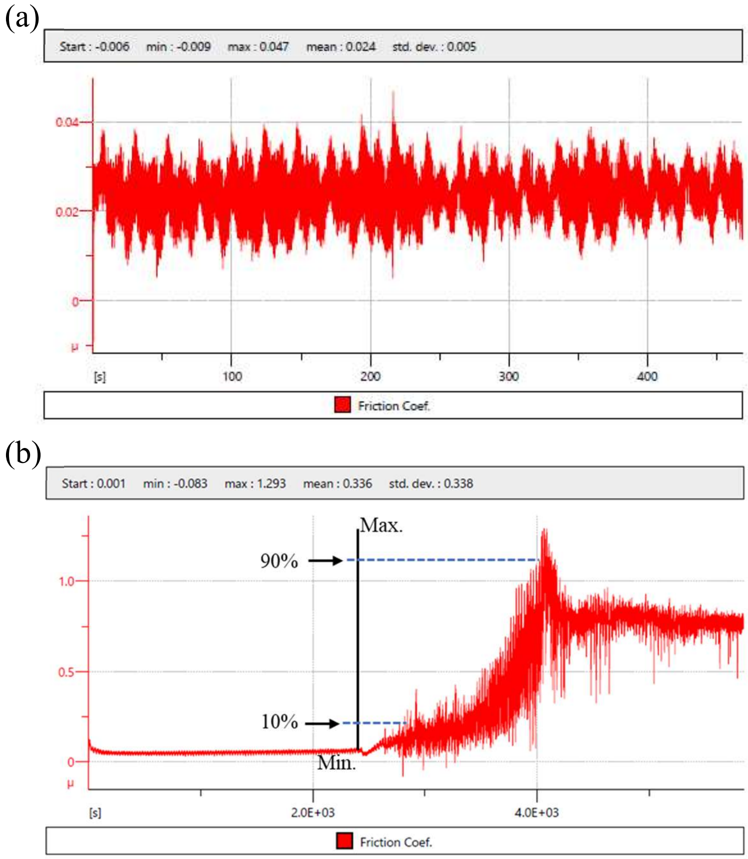


**Supplemental Figure 1.** Representative curves for SBMA hydrogels measured using tribometry for (a) fully hydrated and (b) exposed to ambient conditions samples. (b) indicates the general trend of how T10 and T90 values were calculated.

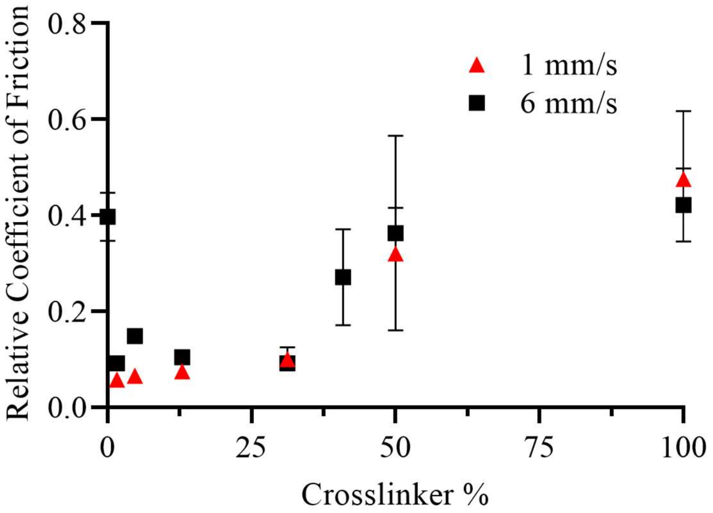


**Supplemental Figure 2.** Relative coefficient of friction for SBMA hydrogels as a function of crosslink density for two probe speeds.

Videos:

Representative videos for the insertion of uncoated (Uncoated_Array) and coated arrays (Coated_Array) into explanted cochleae for force measurements.
