## Supplementary material for "Photografted zwitterionic hydrogel coating durability for reduced foreign body response to cochlear implants": Original Supplemental

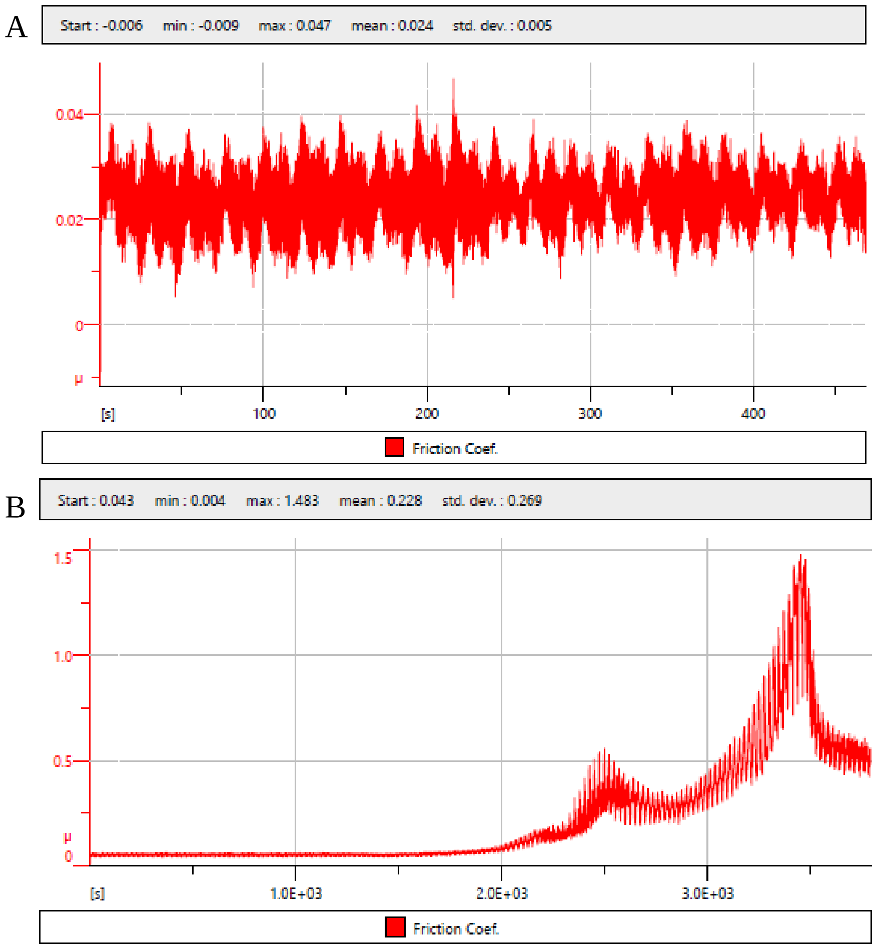


Supplemental Figure 1. Representative curves for tribometry tests where the coefficient of friction is a function of time (seconds) for A) immersed samples and B) samples tested under dry conditions.

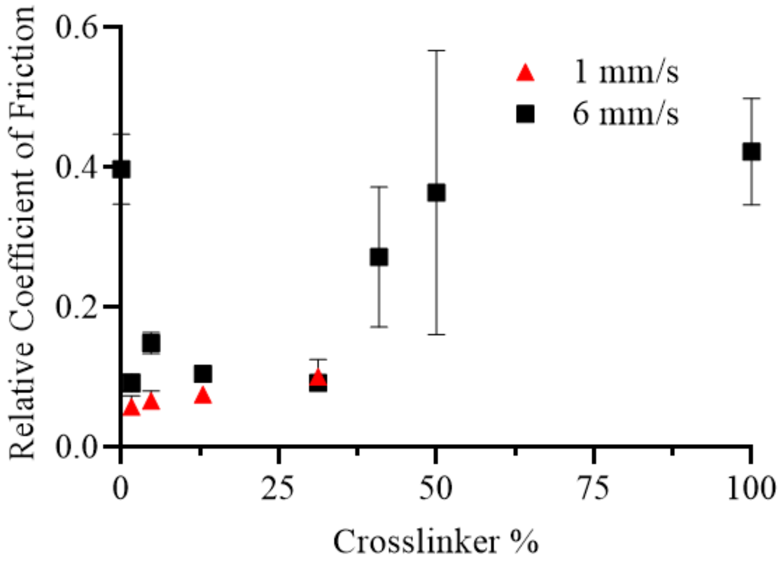


Supplemental Figure 2. The average coefficient of friction (relative to uncoated PDMS) for SBMA-coated samples at two different rotational speeds.
